# Multiplex CRISPR strategy targeting viral genome for agriculture and clinical use

**DOI:** 10.1101/2022.06.09.495443

**Authors:** Zezhong Zheng, Lei Xu, Hongwei Dou, Yixuan Zhou, Xu Feng, Xiangjun He, Zhen Tian, Lingling Song, Yangbin Gao, Guolong Mo, Jiapan Hu, Hongye Zhao, Hongjiang Wei, George M. Church, Luhan Yang

**Affiliations:** Qihan Biotechnology, Hangzhou, China; South China Agricultural University, China; Yunan Agriculture University, Kunming, China; Harvard University, Cambridge, USA

**Keywords:** African Swine Fever Virus, CRISPR, Pig, Agriculture, Xenotransplantation

## Abstract

African Swine Fever (ASF) is a viral disease with exceptionally high lethality in domestic pigs and wild boar worldwide^1, 2^, without any effective vaccine or drug to prevent its spread. In this study, we established a multiplexable CRISPR-Cas-gRNA system targeting 13 genomic loci in the ASF virus genome that could prevent viral replication by cutting its genome. Furthermore, we generated pig strains expressing the multiplexable CRISPR-Cas-gRNA *via* germline genome editing and demonstrated that the gene-edited pigs were more resistant to ASFV infection and less likely to spread the virus upon infection. As far as we know, our study presents the first living organism generated *via* germline editing to demonstrate resistance to viral infection via CRISPR-Cas. We anticipate our work to be helpful for both agricultural and biomedical applications, such as xenotransplantation.

---

African Swine Fever (ASF) is a viral disease with near 100 percent mortality in domestic pigs and wild boar worldwide^1, 2^. The basic reproductive number (R0) has been estimated to be as high as 18.0 for transmission amongst domestic pigs^3^. Over the last 4 years, the outbreak has been reported in 35 countries worldwide, posing a global threat to the swine industry^4, 5^. It is estimated that in 2019 alone, around 50 million pigs died as a consequence of the ASF outbreak with the total economic loss approaching US$111.2 billion^3^. Numerous efforts have been dedicated to developing a safe and effective vaccine against ASF, with only limited success and the potential biosecurity concern of creating new virus variants that may be more deadly and infectious^6^. The current control measures worldwide rely on strict quarantine and stamping out pigs potentially exposed to ASF Virus (ASFV)^2^.

Here, we attempted to engineer an ASF-resistant pig strain using a multiplexable CRISPR-Cas system. ASFV is a double-stranded DNA virus with a genome size of 170 to 190 kbps, encoding 150-200 proteins, depending on the strain^7^. We aimed to specifically cut and destroy the ASFV genome within the pig cells using CRISPR-Cas in order to attenuate ASF replication and prevent viral spread amongst the pig herds.

To this end, we first identified 6 guide RNAs (gRNAs), targeting 13 places in the virus genome (Fig. 1a and Supplementary Table 1). The gRNA-targeting sites were chosen to avoid non-specific cutting into the pig genome (Supplementary Table 1). Next, we designed single transcription constructs to express these 6 gRNAs driven by Type II and Type III RNA polymerase promoters, respectively (Fig. 1c, EF-1a and U6 promoters). We expected the single transcript to be resolved into individual gRNAs via ribozyme sites (Fig. 1a). Within the life cycle of ASFV, it is unclear when and where virus genomic DNA will be targeted by the CRISPR-Cas system^8, 9^ (Fig. 1b). As such, we designed Cas9-expressing constructs with or without nuclear localization signal (NLS). The CRISPR-Cas-gRNA system was packed into a single PiggyBac transposon system (Fig. 1c).

**Fig. 1.**
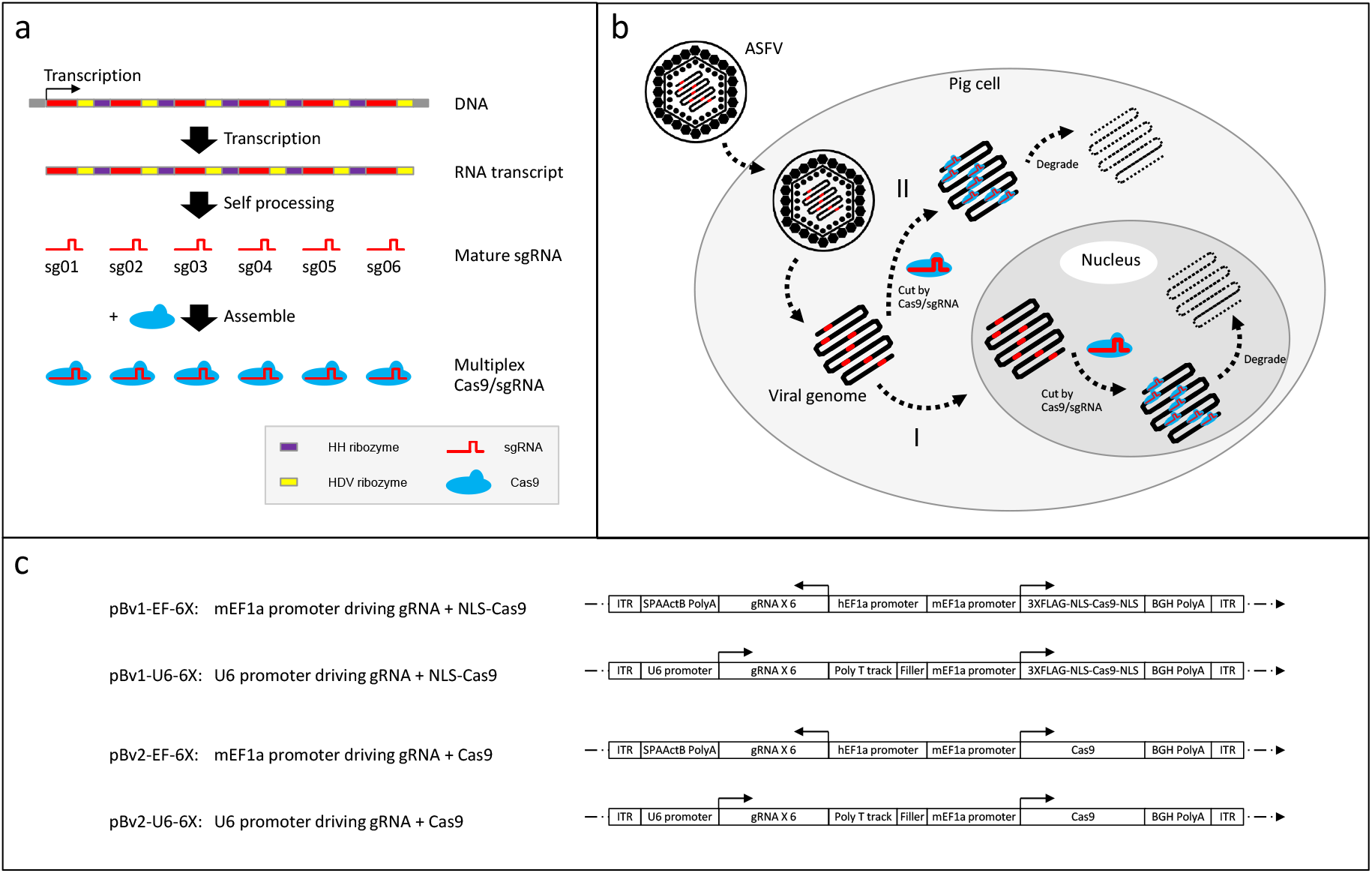
Multiplex CRISPR strategy targeting ASFV genome to protect pig cells from ASFV infection. **a,** DNA construct encoding 6 sgRNAs with each flanked by HH ribozyme and HDV ribozyme can be transcribed under one single promoter into a single RNA transcript which can be automatically processed into 6 mature sgRNA. The mature sgRNA assemble with Cas9 protein to form the catalytic active CRISPR complex. **b,** sgRNA and Cas9 are constitutively expressed in the pig cells, ready to cut invading ASFV genome either in nucleus (route I) or cytoplasm (route II). The viral genome is subsequently degraded and virus replication is stopped or attenuated. **c,** Design illustration of the constructs for expressing both Cas9 protein and 6 sgRNAs. The transgenes are flanked by ITR sequences and inserted into pig genome via Piggybac transposase. For Cas9 expression, either Cas9 with NLS signal or without NLS signal is expressed under mouse EF1a (mEF1a) promoter. For 6 sgRNA expression, either human EF1a (hEF1a) promoter or human U6 promoter is used.

Next, we investigated the cutting efficacy of the CRISPR-Cas-gRNA system on the ASFV genome. Single gRNA transcripts were generated via *in vitro* transcription and its cutting activities were tested against ASFV genomic sequences (Supplementary Fig. 2 and Supplementary Table. 2). All tested gRNAs (5/5) demonstrated accurate and effective cutting activities (Fig. 2a). Later, we conducted genome editing and integrated the CRISPR-Cas-gRNA constructs into the genome of COS-7, a cell line permissive to ASFV infection. Single-cell clones with CRISPR-Cas-gRNA integrated into the genome were isolated, and corresponding Cas9 expressions were confirmed by RT-qPCR (Fig.2b). To examine if the CRISPR-Cas system can protect the cell lines from ASFV infection, we inoculated ASFV into the COS7 clones and monitored the ASFV titer over 5 days. Interestingly, we found that i) the ASFV was significant suppressed at clones where Cas9 expression level is high (Fig. 2b, GC49, GE22, GE64, GF16, GF29 clones) and *vice versa* (FZ2, FY0, FY2, GC7 clones). This suggests that CRISPR-Cas-gRNA can effectively protect COS7 from ASFV infection, the protection level of which is correlated with the expression level of Cas9 (Fig 2c). ii) Both GE and GF clones suppressed ASFV replications, suggesting that either Type II or Type III RNA polymerase was sufficient to drive the multiplexable gRNA expression. iii) Clones with and without NLS completely inhibited ASFV replication (Fig. 2, GE22, GF64 with NLS, GC49 without NLS), indicating that the effect of Cas9 on the ASFV genome was independent of the NLS signal. This observation is consistent with previous reports that ASFV utilizes both the host nucleus and cytoplasmic “virus factory” to replicate its DNA^10, 11^, so that its genome is exposed in both sites. Taken together, the data demonstrates that CRISPR-Cas9-gRNA can effectively prevent ASFV replication *in vitro* by cutting the genome at multiple locations.

**Fig. 2.**
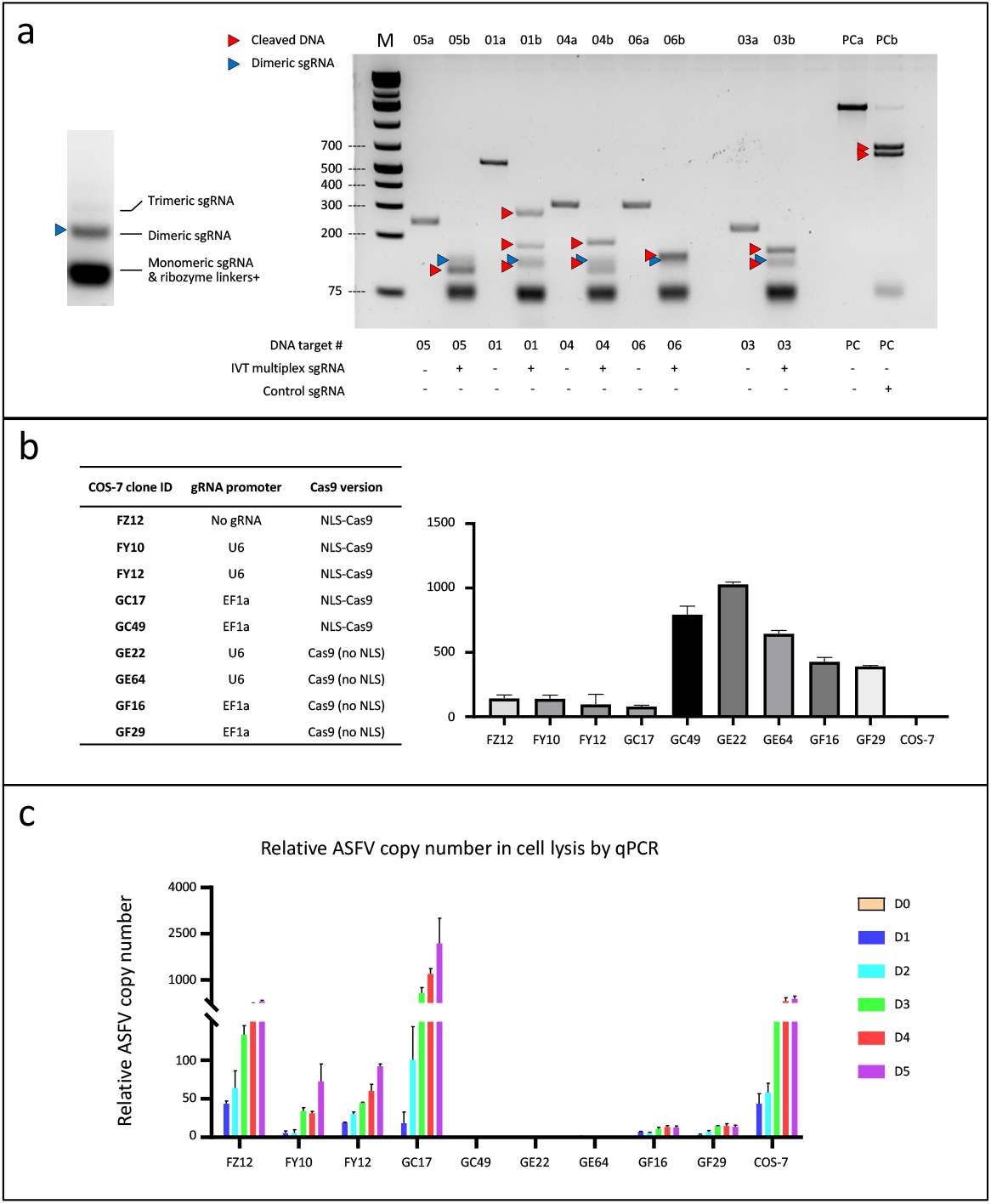
Multiplex CRISPR strategy can efficiently cut ASFV DNA in vitro and different construct design has different effects on preventing ASFV replication in COS-7 cells. **a,** Left panel, the single RNA transcript generated by in vitro transcription (IVT) can be efficiently processed into monomeric gRNA with few dimeric (blue triangle) and trimeric gRNA. Right panel, the IVT single RNA transcript can mediate efficient cleavage of PCR amplicons amplified from the ASFV genome in *in vitro* Cas9 cleavage assay. “DNA target #” indicates the PCR amplicon corresponding to the respective gRNA. The starting PCR amplicons (lane 01a, 03a, 04a, 05a, 06a, PCa) are digested into 2 or more fragments (red triangles, lane 01b, 03b, 04b, 05b, 06b, PCb. PC: positive control) after co-incubation with IVT single RNA transcript and Cas9 protein. **b,** Engineered COS-7 single-cell clones with different gRNA promoters and Cas9 versions (left table) and relative Cas9 expression levels in those clones (right panel) determined by RT-qPCR. **c,** ASFV replication in different COS-7 single-cell clones measured by ASFV copy number in cell lysis using qPCR over 5 days. Strong inhibition of ASFV replication is only observed in COS-7 clones with high levels of Cas9 expression (GC49, GE22, GE64).

Having validated the ASFV-resistant activity of the CRISPR-Cas9-gRNA system *in vitro,* our next goal was to germline engineer a pig strain resistant to ASFV infection by integrating an effective construct into the pig genome. We chose the NLS-Cas9-EF1a-gRNA construct (identical to the construct of the GC49 COS7 clone) as it demonstrated significant ASFV suppression *in vitro* (Fig. 2c). To this end, we first integrated the construct via the PiggyBac transposon system into the fibroblasts of a Large White pig strain. After isolating the single cells clones with the appropriate genetic modification, we performed pig cloning *via* somatic cell nuclear transfer technology (SCNT) and successfully obtained pigs with intended modifications (Fig. 3b). In addition, we validated the genomic integration and the expression level of Cas9 in tissue samples obtained from our engineered pigs (Fig. 3a, GI58 P10, GI58 P12).

**Fig. 3.**
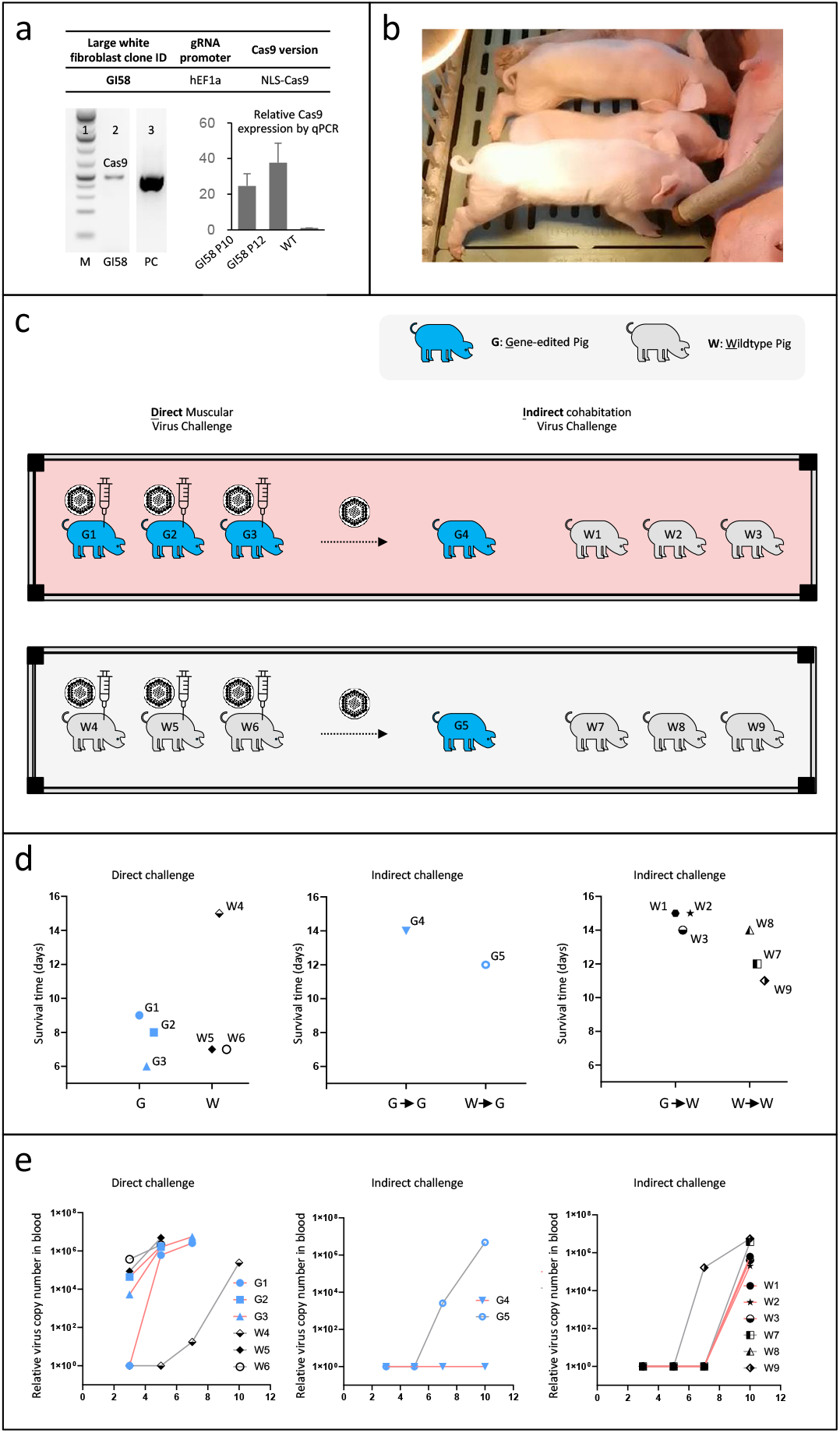
Production of transgenic pigs constitutively expressing Cas9 and 6 sgRNA targeting ASFV genome and virus challenge of the transgenic pigs. **a,** Top table, the large white pig fibroblast single cell clone GI58 harbors the pBv1-EF-6X construct with hEF1a promoter driving sgRNA and mEF1a promoter driving Cas9 with NLS. Bottom left panel, presence of the transgene cassette was confirmed by PCR for Cas9 gene in genomic DNA of GI58. Bottom right panel, Cas9 expression was confirmed by RT-qPCR in the fibroblasts of the cloned pigs (GI58P10 and GI58P12) generated from GI58 fibroblasts. **b,** photo of the cloned pigs generated from GI58 fibroblasts. **c,** ASFV virus challenge design of the gene-edited pigs and wildtype pigs. Direct virus challenge was performed by muscular injection, while indirect virus challenge was performed by cohabitation of the pigs in the same room with pigs under direct virus challenge. Each rectangle with four black squares indicates a separate room. In light-red room 3 gene-edited pigs were subjected to direct virus challenge, while in the light-grey room 3 wildtype pigs were subjected to direct virus challenge. The additional 1 gene-edited pig and 3 wildtype pigs in each room are subjected to indirect virus challenge. **d,** Survival of the pigs in direct and indirect virus challenge. No statistical difference in survival time was observed in the two virus challenge modes comparing gene-edited pigs with wildtype pigs under the same condition. The arrow indicates indirect virus challenge, e.g., W->G means wildtype pigs were under direct virus challenge and gene-edited pigs were under indirect virus challenge released by infected wildtype pigs. **e,** ASFV titer in blood samples of the pigs in direct and indirect virus challenge measured by qPCR. Gene-edited pigs showed no detectable virus titer, much lower than wildtype pigs, in indirect virus challenge released by infected gene-edited pigs (middle panel). No difference was observed in other conditions.

We investigated if the established pig strain with multiplexable CRISPR-Cas-gRNA modification was resistant to ASFV infection. To this end, transgenic and wild-type pigs were challenged directly or indirectly with ASFV (Fig. 3c). In the direct-challenge group, the same amount of concentrated ASFV was injected into the muscle of transgenic pigs (3 pigs: G1, G2, G3) and wild-type pigs (3 pigs: W4, W5, W6). In addition, to evaluate the capacity of pigs to resist ASFV horizontal transmission, we kept one transgenic pig and three wild-type pigs co-housed with the direct-challenge groups as the indirect-challenge groups (G4, W1, W2, W3 co-housed with G1-G3; G5, W7, W8, W9 co-housed with W4-W6). Viral titers in the pig blood and survival time were closely monitored. Consistent with the previous reports^12, 13^, intramuscular injection of ASFV was lethal and wild-type pigs died within 7 days, although surprisingly, one outlier (W4) survived over 2 weeks with elevated blood ASFV titer (Fig 3e). The wild-type pigs who had gone through indirect challenges were dead with a delayed time course (Day 10-Day 14). CRISPR-Cas9-gRNA germline modification could not prevent the pigs from death but did slow the progression of diseases in our experiment. i) We observed that transgenic pigs (G1-G3) survived longer (~2-3 days) upon direct ASFV challenges (Fig 3d) and with a late onsite increase of viral titer in the blood (Fig. 3e). ii) we found the consistent trend that pigs co-housed with infected transgenic pigs have a longer survival time compared with pigs co-housed with infected wild-type pigs (Fig. 3d G4 survived longer than G5, W1-W3 survived longer than W7-W9 on average). This data was also corroborated by the difference in viral titer curves detected in these pigs (Fig. 3e). Together, our data suggested that CRISPR-Cas9-gRNA in the genome could lower ASFV viral titer and delay the virus from spreading. More interestingly, we did not detect any virus in the blood of G4, a transgenic pig co-housed with the infected transgenic pigs (G1-G3); this pig also exhibited the best survival and health status until the end of this study, suggesting that pigs harboring CRISPR-Cas9-gRNA modification acquired some degrees of immunity against the ASFV infection from other pigs.

In summary, we established a multiplexable CRISPR-Cas-gRNA system targeting 13 genomic loci in the ASFV genome, which attenuated viral viability by cutting its genome. We demonstrated that Cas9-gRNA could effectively target the ASF genome and control the viral titer in the COS7 system. Furthermore, we generated pig strains expressing the multiplexable CRISPR-Cas-gRNA *via* germline genome editing. We demonstrated that the gene-edited pigs were more resistant to ASFV infection and less likely to spread the virus upon infection.

As far as we know, our study presents the first living organism generated *via* germline editing to demonstrate resistance to viral infection via CRISPR-Cas. Previous efforts targeting one site ASFV genome via CRISPR-Cas-gRNA in the cell line, demonstrated ASFV replication can be successfully suppressed by manipulating the CP204L gene alone^14^. However, it remained unknown whether such an *in vitro* effect could be translated into *in vivo* protection. In addition, targeting a single genomic locus is vulnerable to escape mutants. A multiplexable gene editing approach is thereby essential to eliminate the virus before accumulating mutants circumvents all the host’s defense mechanisms.

Notably, we see more remarkable effects in the *in vitro* setting than that *in vivo*, which might be attributed to different viral loads between the cell line experiments and the *in vivo* pig experiments. Intramuscular injection exposes the pig to orders of magnitude higher viral titers than typical infection through contacts. Encouragingly, G4, a transgenic pig co-housed with the transgenic pigs with ASFV injection, survived until the end of the study. G4 may benefit from more resistance to ASFV and less exposure to ASFV shedding from infected transgenic pigs. We look forward to future experiments with bigger sample size to quantify the protection towards ASF on the population level.

In the future, more advanced genome-engineering work can be done to enhance herd immunity against ASFV infection, including increasing the Cas9 and gRNA expression level (Fig. 2b), delivering both Cas9 constructs with and without NLS so that it could repel the virus throughout the life cycle (Fig 1b and 2c). Moreover, the construct could be engineered to conditionally express upon infection to avoid potential interference with normal cell physiology.

The prevention and control of ASFV are not only crucial to agricultural applications but also have significant clinical implications. Recent clinical advancement in xenotransplantation encourages more clinical testing of transplanting pig organs into human patients^15, 16^; however, it also potentiates the risk of novel zoonotic diseases by either endogenous origin such as PERVs^17, 18^ or by external source such as ASFV and CMV. In the wake of the COVID pandemic, preempting any possibility of horizontal animal virus transmission through pig organs is critical, especially considering that ASFV is widespread and affects pig industry globally^19^. In addition, our approach can be utilized to engineer pig organs with novel biological functions, such as targeting HBV for liver cancer patients who need a liver transplant.

## Acknowledgements

We thank G. Yang of Harvard University for reading our manuscript. The Animal Biosafety Level-3 Laboratory (ABSL-3) was co-provided by the Jinyu Bio. And we are grateful to colleagues at Qihan Bio for their technical assistance and discussions.

## Author contributions

L.Y., G.M.C., H.W. and H.Z. envisioned and supervised the whole project, H.W. supervised pig cloning and production. Y.G. and Z.Z. designed the experiments. L.X., H.D., Y.Z., Z.T., and L.S. performed experiments. X.F. and X.H. analyzed the data. G.M., L.X. and J.H. wrote and revised the manuscript.

## Additional information

Supplementary information as submitted

## Methods

### CRISPR-Cas9 gRNA design

R library DECIPHER was used to design specific gRNAs (Supplementary Table. 2)

### Plasmid construction

pBv1-EF-6X, pBv1-U6-6X, pBv2-EF-6X and pBv2-U6-6X (Fig. 1c) were constructed by Golden Gate Assembly (NEB Golden Gate Enzyme Mix (BsaI-HF2), New England BioLabs) with 1 of the 4 backbone plasmids pBv1-EF-BB / pBv1-U6-BB / pBv2-EF-BB / pBv2-U6-BB and all the 6 TOPO-RGR (RGR: ribozyme-sgRNA-ribozyme) plasmids following manufacturer’s instructions. The backbone plasmids were constructed by Gibson Assembly and NLS-Cas9/Cas9 was PCR amplified from pX330-U6-Chimeric_BB-CBh-hSpCas9 vector^20^. Each of the TOPO-RGR fragment was constructed by overlapping PCR followed by TOPO cloning using pClone007 Versatile Simple Vector Kit (Tsingke). The sequences of the multiplex sgRNA designs using either RNA pol II mEF1a promoter and U6 promoter are illustrated in Supplementary Fig 1.

### In vitro transcription (IVT) and Cas9 cleavage assay

PCR products were amplified from pBv1-U6-BB plasmid with primers:

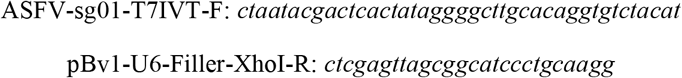

to append the T7 promoter at the 5’ end of the multiplex guide RNA cassette. The PCR products were TOPO cloned using pClone007 Versatile Simple Vector Kit (Tsingke) and the TOPO plasmid were isolated. The TOPO plasmid containing the multiplex guide RNA cassette was digested with *Xho* I and purified to be used as IVT template. IVT was carried out using MEGAscript T7 Kit (Thermo Fisher Scientific) and purified by MEGAclear Kit (Thermo Fisher Scientific) following manufacturer’s instructions. Purified IVT product was analyzed using 2% agarose gel electrophoresis (Fig. 2a). DNA substrates containing the guide RNA target sites were PCR amplified from purified ASFV genomic DNA using primers detailed in Supplementary Table 2. In vitro Cas9 cleavage assay was carried out using purified IVT product and Cas9 protein (Cas9 Nuclease, S. pyogenes, New England BioLabs) following manufacturer’s instructions. The digested products were analyzed using 2% agarose gel electrophoresis (Fig. 2a).

### Cell culture of pig fibroblast and COS-7 cells

Pig fibroblast cells were isolated from wildtype large white ear sample and cultured in Dulbecco’s modified Eagle’s medium (DMEM, high glucose, GlutaMAX Supplement, pyruvate, Thermo Fisher Scientific) supplemented with 20% fetal bovine serum (Thermo Fisher Scientific), 1% Penicillin/Streptomycin (Pen/Strep, Thermo Fisher Scientific) and 1% HEPES (Sangon). Pig fibroblast cells were maintained in a humidified tri-gas incubator at 37°C and 5% CO_2_, 5% O_2_ and 90% N_2_.

COS-7 cells were cultured in Dulbecco’s modified Eagle’s medium (DMEM, high glucose, GlutaMAX Supplement, pyruvate, Thermo Fisher Scientific) supplemented with 10% fetal bovine serum (Thermo Fisher Scientific), 1% Penicillin/Streptomycin (Pen/Strep, Thermo Fisher Scientific) and 1% HEPES (Sangon). COS-7 cells were maintained in a humidified incubator at 37°C with 5% CO_2_.

### Gene editing and single cell cloning of the pig fibroblast and COS-7 cells

Gene editing of COS-7 cells were performed by electroporation of COS-7 cells with Super PiggyBac plasmid (PB210PA-1, System Biosciences) and one of the 5 plasmids of pBv1-EF-6X, pBv1-U6-6X, pBv2-EF-6X, pBv2-U6-6X and pBv1-U6-BB plasmids using Neon Transfection System (Invitrogen) under condition of 1050 V, 30 ms, 2 pulses. The transfected cells were single-cell sorted using SONY SH800S into 96-well plates, and single cell clones were grown and clones positive for Cas9 gene cassette by PCR were chosen for Cas9 expression analysis. The final clones chosen for analysis are FY10, FY12, GC17, GC49, GE22, GE64, GF16, GF29 and FZ12 (Fig. 1b).

Gene editing of pig fibroblast cells were performed by electroporation of wildtype large white pig fibroblast cells with a plasmid encoding Piggybac transposase and pBv1-EF-6X plasmid using Neon Transfection System (Invitrogen) under condition of 1450 V, 10 ms, 3 pulses. The edited cells were single-cell sorted using SONY SH800S into 96-well plates, and single cell clones were grown and clones positive for Cas9 gene cassette were chosen for Cas9 expression analysis. Clone number GI58 with expression of Cas9 was chosen for pig cloning (Fig. 3a).

### Virus challenge in COS-7 cells in vitro

Virus challenge experiments were standardized according to the Laboratory Biosafety Manual and strictly performed in the P3 biosafety laboratory. The COS-7 cell virus challenge experiments were carried out in the Animal Biosafety Level-3 Laboratory (ABSL-3) at South China Agricultural University (Guangzhou, China). Wildtype COS-7 cells were used to adapt ASFV to more robust replication after passing ASFV in COS-7 cells for a few generations. Supernatant containing adapted ASFV virus was harvested from the infected COS-7 at P9 adaption and was used for in vitro virus challenge. Gene-edited COS-7 single-cell clones were seeded into 24-well plate at 2×10^5^ cells/well, 12 wells/clone, and grown in regular media for 1 day before changed into 500 μl media containing Dulbecco’s modified Eagle’s medium supplemented with 2% fetal bovine serum, 1% Penicillin/Streptomycin and 1% HEPES. 10 μl of the adapted virus was inoculated into each well of the COS-7 single cell clones, and two wells were harvested each day and the cells were lysed for further analysis.

### PCR, qPCR and RT-qPCR

PCR for the presence of Cas9 transgene was performed using primers:

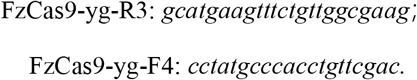

qPCR for ASFV quantification was performed with primers:

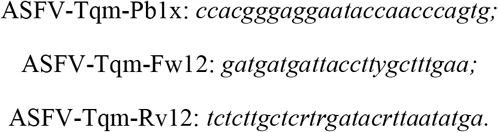

RT-qPCR for quantification of Cas9 RNA expression level was performed using primers:

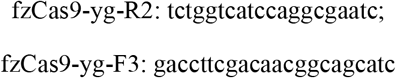

with cDNA reverse-transcribed from isolated total RNA from corresponding cells.

### Somatic cell microinjection to produce SCNT embryos and embryo transfer for pig cloning

Animal experiments involving ASFV were standardized according to the Laboratory Biosafety Manual and strictly performed in the P3 biosafety laboratory. The animal experiments were carried out in the Animal Biosafety Level-3 Laboratory (ABSL-3) of the Spirit Jinyu Biological Pharmaceutical Co., LTD (Neimenggu, China). The somatic cell microinjection procedure and SCNT was conducted as previously described^19, 22-24^.

### Virus challenge in gene-edited pigs in vivo

17 animals were enrolled in the experiment, which included 5 transgenic pigs and 12 wild-type pigs. 14 animals were set into two groups and challenged with ASFV genotype II strain GZ201801. Group setting and challenge route were shown in the following table.

**Experimental animal grouping and challenge**
| **Groups** | **Number of animals (head)** | **Challenge route** | **Challenge dosage** |
| --- | --- | --- | --- |
| **Challenge group of transgenic pigs** | **3 (transgenic pigs challenge) +3 (wild-type pigs cohabit) +1 (transgenic pigs cohabit)** | **Intramuscular injection** | **10<sup>3</sup>HAD<sub>50</sub>/mL**<br>**2 mL** |
| **Challenge group of wild-type pigs** | **3 (wild-type pigs challenge) +3 (wild-type pigs cohabit) +1 (transgenic pigs cohabit)** | **Intramuscular injection** | **10<sup>3</sup>HAD<sub>50</sub>/mL**<br>**2 mL** |
| **Wild-type pigs (negative)** | **3** | **—** | **—** |

Note: Due to the limited number of transgenic pigs, they were arranged in to two groups to test the resistance to direct challenge and cohabit challenge.

The challenge experimental groups and the non-challenge negative group were fed by different workers and housed in isolated rooms to prevent cross-infection.

Before the experiment, all animals were ear-tagged and tested negative for ASFV antigens and antibodies. On day-1 of the experiment, EDTA anticoagulated blood and whole blood, as well as oropharyngeal and anal swabs were collected from all animals. Clinical symptom scores and temperature (rectal) of animals were recorded daily. Clinical symptoms were evaluated by the caretakers according to the flowing table.

**Clinical symptom index scoring criteria**
| **Observation item** | **Scoring criteria** | **Score** |
| --- | --- | --- |
| **a Appetite** | **Normal** | **0** |
|  | **Slight loss** | **1** |
|  | **Significant loss** | **2** |
|  | **Complete loss** | **3** |
| **b Mental state** | **Normal** | **0** |
|  | **Slow movement and low spirits** | **1** |
|  | **Often get down and depressed** | **2** |
|  | **Lie on the ground and drowsy** | **3** |
| **c Temperature (rectal temperature)** | **Normal (38°C—40°C)** | **0** |
|  | **Low fever (40°C—42°C)** | **1** |
|  | **High fever (> 42°C)** | **3** |
| **Overall clinical score = a + b + c** |  |  |

Animals were challenged on day-4. For all experimental animals, oropharyngeal swabs and anal swabs were collected daily from day-4 of the experiment. Besides, EDTA anticoagulated blood and whole blood were collected every two days from day-1 until the end of the experiment or the death of animals. During the experiment, dying animals were euthanized and dissected for sampling. The experiment ended on day-18. All remaining animals were euthanized and then dissected for sampling.

ASFV antigens were tested in EDTA anticoagulated blood samples and swab samples through real-time PCR. Serum samples were tested for ASFV antibodies.

Fresh and fixed tissues of the heart, liver, spleen, kidney, lung, mandibular lymph nodes, thoracic lymph nodes, and mesenteric lymph nodes were collected in the necropsy. ASFV antigen and mRNA of IL-1β, IL-6, IL-8, and TNF-α were tested in the fresh tissue tissues through real-time PCR. Histopathology changes were evaluated in the fixed tissues. Immunohistochemistry tests were also performed on the fixed tissue to analyze the virus distribution in the organ systems.

**Supplementary Table 1.**
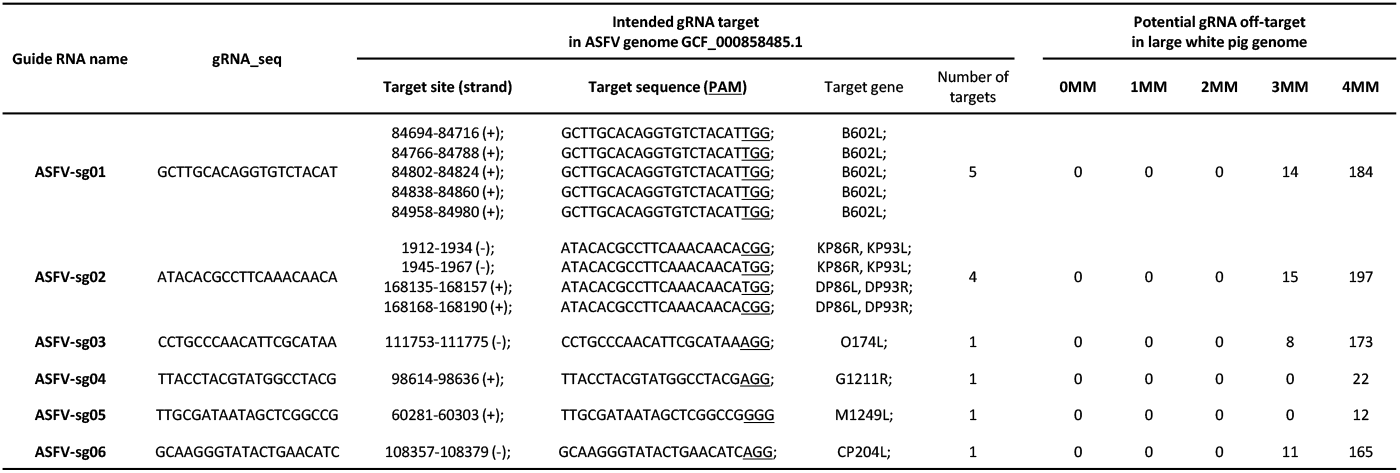
Intended guide RNA targets in ASFV genome and potential off-targets in pig genome. Six guide RNAs, ASFV-sg01-06, are designed to target ASFV genome based on BA71V strain (RefSeq: GCF_000858485.1). ASFV-sg01 targets the viral genome 5 times, and ASFV-sg02 targets the viral genome 4 times. ASFV-sg03-06 targets the viral genome 1 time. The potential gRNA off-targets in large white pig genome are predicted bioinformatically. No predicted off-targets for all 6 guide RNAs even at relaxed criteria of 2 mismatches (MM). Although unlikely to be true off-target sites, potential off-targets with 3 or 4 MM are also listed.

**Supplementary Fig. 1.**
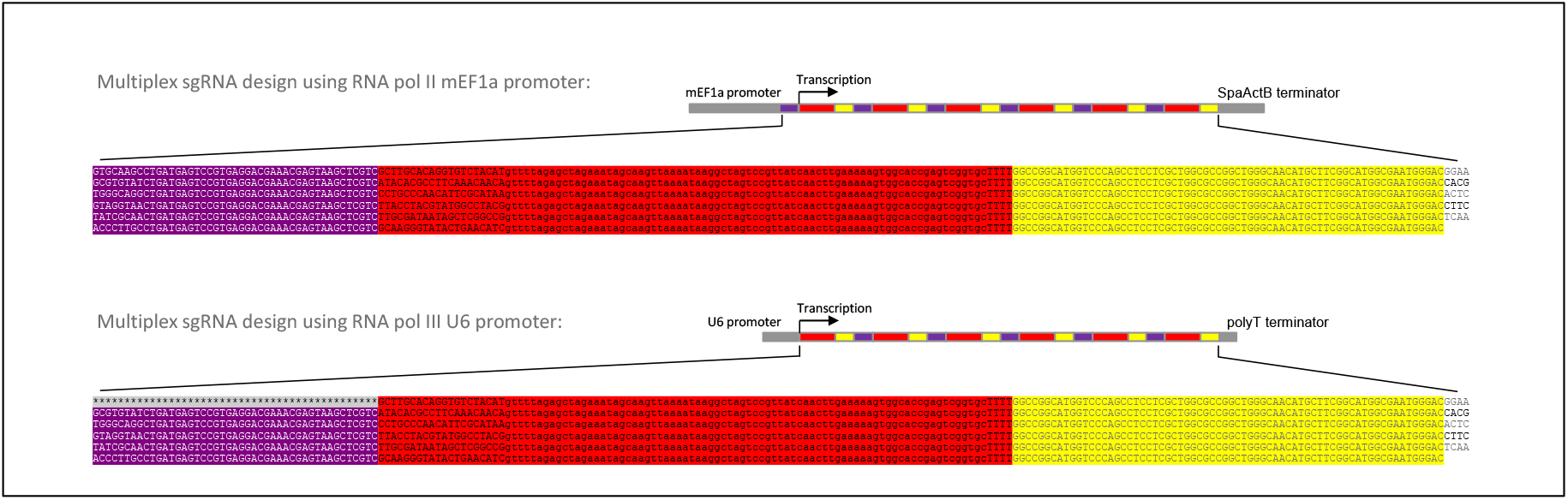
Sequence of the multiplex sgRNA design using mEF1a promoter or U6 promoter. Purple color, HH ribozyme. Red color, sgRNA. Yellow color, HDV ribozyme. Top panel, multiplex sgRNA design using RNA pol II mEF1a promoter. Each sgRNA sequence is flanked by HH ribozyme and HDV ribozyme. Bottom panel, multiplex sgRNA design using RNA pol III U6 promoter. Compared with multiplex sgRNA design using mEF1a promoter, the first sgRNA does not have HH ribozyme at its 5’ end.

**Supplementary Fig. 2.**
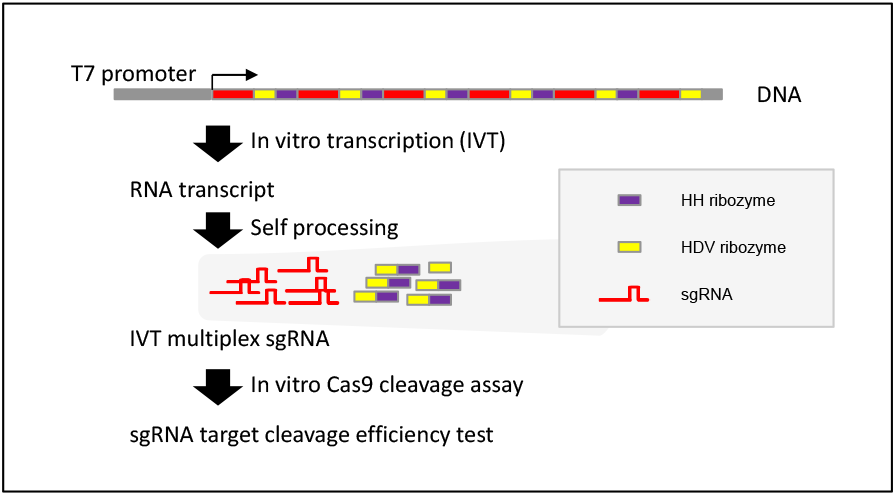
Multiplex guide RNA efficiency test using in vitro Cas9 cleavage assay. A T7 promoter is added at the 5’ end of the multiplex sgRNA construct similar to the U6 promoter design (Supplementary Fig. 1). In vitro transcription (IVT) using T7 RNA polymerase produces RNA transcripts which can be automatically processed into functional individual sgRNAs and ribozyme linker byproducts. The cleavage efficiency of the individual sgRNAs are further tested in in vitro Cas9 cleavage assay using PCR amplicons containing the sgRNA target sites amplified from ASFV genome.

**Supplementary Table 2.**
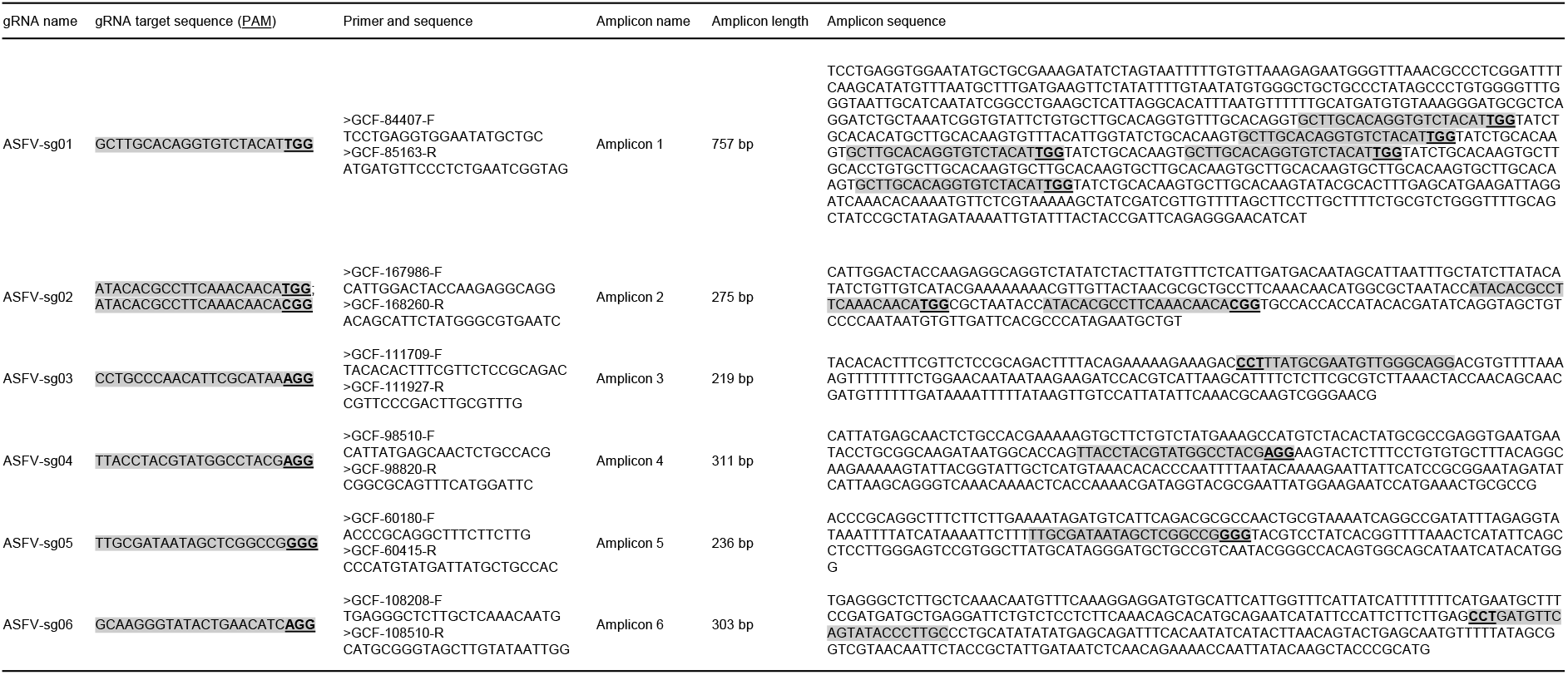
Primers and amplicons used in in vitro Cas9 cleavage assay. DNA targets of the six guide RNAs ASFV-sg01-06 are amplified from ASFV genomic DNA using the primer pairs listed. The guide RNA targets in the PCR amplicons are shaded in grey and the PAM sites are underlined. Predicted cleavage patterns: Amplicon 1 = 757 bp = (303 + 72 + 36 + 36 + 120 + 190) bp. Amplicon 2 = 275 bp = (165 + 33 + 77) bp. Amplicon 3 = 219 bp = (60 + 159) bp. Amplicon 4 = 311 bp = (120 + 191) bp. Amplicon 5 = 236 = (117 + 119) bp. Amplicon 6 = 303 bp = (165 + 138) bp.

